# Highly efficient multiplex genome editing in dicots using improved CRISPR/Cas systems

**DOI:** 10.1101/2022.01.10.475635

**Authors:** Bingjie Li, Yun shang, Lixianqiu Wang, Jing Lv, Fengjiao Wang, Jiangtao Chao, Jingjing Mao, Anming Ding, Xinru Wu, Mengmeng Cui, Yuhe Sun, Changbo Dai

## Abstract

CRISPR/Cas9-mediated gene editing provides a powerful tool for dissecting gene function and improving important traits in crops. However, there are still persisting challenges to obtain high homozygous/bi-allelic (ho/bi) mutations in dicot plants. Here, we develop an improved CRISPR/Cas9 system harboring a *calreticulin-like* gene promoter, which can boost targeted mutations in dicots. Additionally, the pDC45_dsg construct, combining a *35Spro-tRNA_sgRNA-EU* unit and *PCE8pro*-controlled Cas9, can achieve more than 80.0% ho/bi mutations at target sites in allotetraploid tobacco. We construct pDC45_Fast system that can simultaneously fulfill gene editing and shorten the life span of T0 generation tobacco and tomato. This study provides new tools for improving targeted gene mutagenesis in dicots, and makes manipulations of genes in *Solanum* more feasible.

## Background

Genome editing mediated by Clustered regularly interspaced short palindromic repeats (CRISPR)/Cas9 systems are widely used to study functional genomics and improve the agronomic traits in crops [1, 2]. To date, various plant-specific CRISPR/Cas expression systems have been developed and applied in model plants, including Arabidopsis, *Oryza sativa*, and *Nicotiana tabacum*, but editing efficiencies vary widely [3–5]. A prototypical CRISPR/Cas system with the CaMV *35Spro* and polymerase III (pol III) promoter, can accomplish genome editing in many plants, but generates relatively few homozygous/bi-allelic (ho/bi) mutations, especially in *Solanum* species [3]. Almost all research groups achieved less than 36.4% ho/bi editing efficiencies via CRISPR/Cas9 system in allotetraploid tobacco [5–12], thus more work is needed to develop new systems to boost the target mutations in *Solanum* plants. Here, we report improved CRISPR/Cas9 cassettes, harboring a novel type promoter and the tRNA-sgRNA transcript unit, can generate robust editing efficiency in T0 generation tobacco and tomato.

## Results and discussion

Considering that using the proper promoters for CRISPR/Cas systems are critical for gene editing, we presumed that promoters exhibiting high activities in tobacco calli might improve the mutation efficiency of CRISPR/Cas9-mediated genome editing in plants. Therefore, RNA-seq was employed to identify plant callus expression (*PCE*) genes from different tobacco developmental stages. One of these genes, *PCE8*, encoding a calreticulin-like protein, had an extremely high expression level in the early stage of tobacco shoot regeneration. Since the *PCE8* promoter (*PCE8pro*) possess cis-elements related to meristem expression, hormone and wound response etcetera (Supplementary Fig. 1), we speculated that the *PCE8pro* was suitable to produce efficient targeted editing in the tobacco genome. To validate this hypothesis, we developed two types of CRISPR/Cas9 expression cassettes, pDC30 and pDC40, in which the sgRNA is driven by the At*U6-26* promoter [13], and *SpCas9* is separately controlled by the 35S and *PCE8* promoter. Then, we developed pDC45 and pDC45_dsg systems relying *35Spro*, the tRNA-mediated polycistronic gRNA transcript unit [14], and an EU terminator [15] to achieve multiplex genome editing in tobacco (Fig. 1a).

**Fig. 1.**
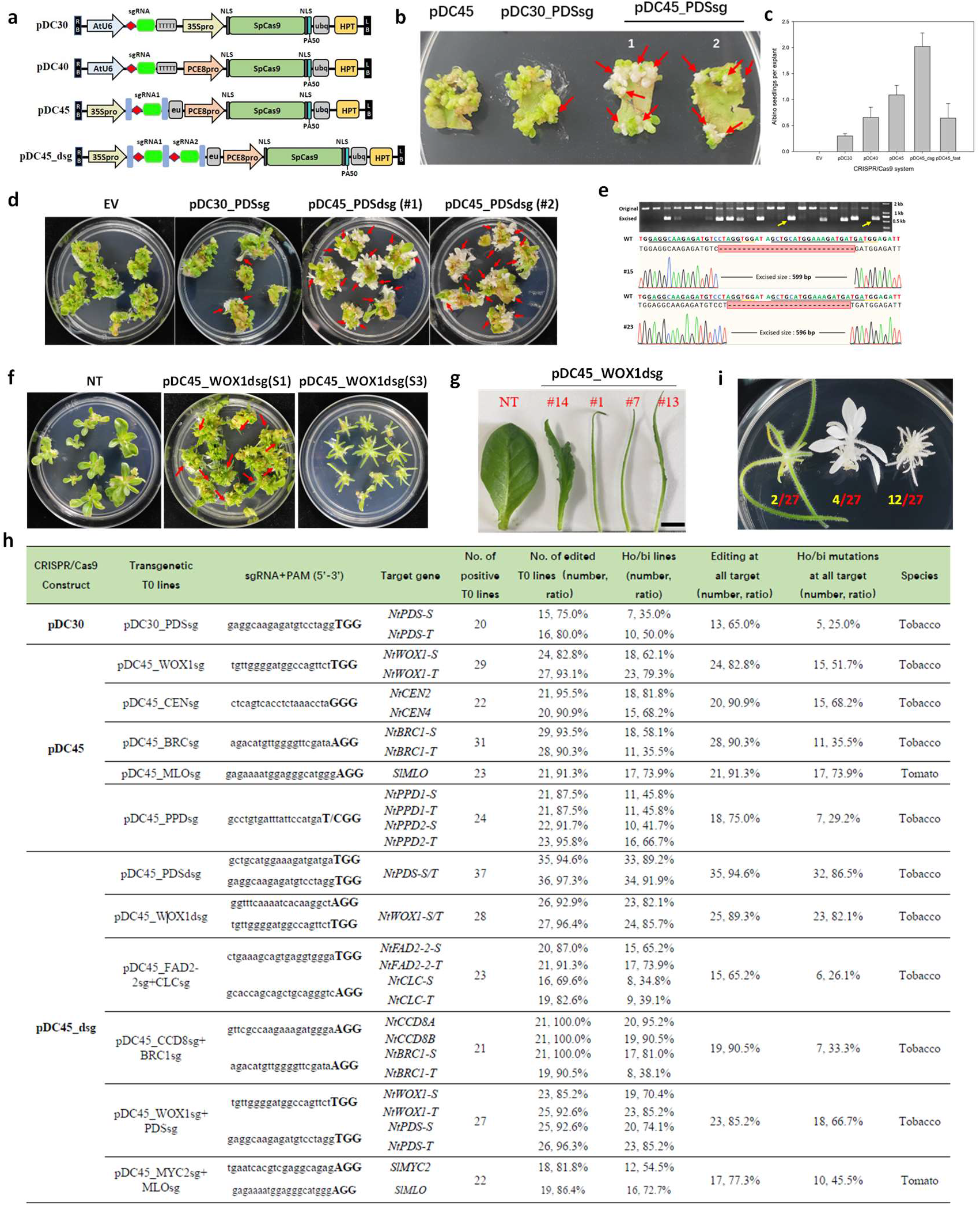
Highly efficiency of gene editing in plants. **a** Schematic illustration of CRISPR/Cas9 systems used in this study. NLS, nuclear location sequence; ubq; *AtUBQ1* terminator; eu, intronless tobacco *extension* terminator; bar in light blue shows tRNA sequence; HPT, *hygromycin phosphotransferase* gene. **b** The albino phenotypes of tobacco T0 seedlings generated via various CRISPR/Cas9 expression cassettes. 1 and 2 show the randomly selected explants at early stage of regeneration. The red arrowheads indicate the albino phenotype of *NtPDS* disruption shoots. EV, empty pDC45 vector, **c** Comparison of the ho/bi mutations based on the ratios of albino plantlets generated by various CRISPR/Cas9 systems, the standard deviation (SD) came from three replicates. **d** The phenotype of transgenic tobacco lines generated by various pDC constructs. #1 and #2 plates show thirty-days explants from two individual transformations. The red arrowheads indicate the absolute albino shoots. **e** Agarose gel electrophoresis and Sanger sequencing of fragments deletions generated by pDC45_PDSdsg. Yellow arrow show the #15 and #23 PCR product selected for Sanger sequencing. **f** Phenotypes of *NtWOX1* mutated plants produced by pDC45_WOX1dsg, NT, seedlings lacking T-DNA insertion, the red arrowheads show lines with bladeless leaf, S1 and S3 mean different stages of tissue regeneration. **g** The leaf phenotype of pDC45_WOX1dsg transformed tobacco, number in red mean individual T0 lines, scale bar, 1 cm inlength. **h** Summary of the genotyping results on stable T0 transgenic plants by various CRISPR/Cas9 systems. **i** The diverse phenotypes of ho/bi T0 lines with pDC45_PDSsg+WOX1sg construct, number in yellow and red show number of ho/bi lines and Hyg-resistance lines, respectively.

Tobacco *PHYTOENE DESATURASE* (*NtPDS*) was selected to investigate the editing efficiency of our systems. Seedlings with disrupted *NtPDS* genes have an albino phenotype, making it easy to determine whether both *NtPDS* alleles are edited in tobacco S- and T-genomes (Fig. 1b). For T0 lines transformed by pDC30 cassette, we observed five to nine absolute albino seedlings among 19 to 27 randomly selected explants from three individual experiments, resulting in an average of 0.30 albino shoots per explant (Fig. 1c). Using the same strategy, we obtained 0.66, 1.09, and 2.02 albino seedlings per explant on average from pDC40, pDC45, and pDC45_dsg transformed tobacco, which are respectively 2.20-, 3.63-, and 6.73-fold higher than in the pDC30 construct (Fig. 1c, d). Additionally, 52.2% of albino plants had fragment deletions from two cleavage sites of *NtPDS* sgRNAs (Fig. 1e), suggesting that *PCE8pro*-controlled CRISPR/Cas9 is highly efficient in generating targeted mutations in tobacco.

The high ho/bi mutations (82.1%) of pDC45_dsg transgenic lines were further confirmed through dual gRNAs-mediated targeting tobacco *WUSCHEL RELATED HOMEOBOX 1*-like genes (*NtWOX1*). Unlike plants without T-DNA construct, many T0 transgenic lines with bladeless leaves were obtained from randomly selected tobacco leaf discs due to disruption of both *NtWOX1* loci (Fig. 1f, g).

To further investigate the editing capability of the pDC45 system in tobacco, we designed gRNAs to target *BRANCHED 1 (NtBRC1), CENTRORADIALIS* (*NtCEN*), and *NtWOX1* alleles separately in both S- and T-genomes. Hi-TOM sequencing [16] results indicated that 82.8% to 95.5% of plants were mutated at each target site, and 82.8% to 90.9% were edited at both target loci. The ho/bi mutations at *NtBRC1-T* alleles had relatively lower editing efficiencies (35.5%) compared to mutations at *NtCEN* (68.2%) and *NtWOX1* (51.7%) sites (Fig. 1h). Although fewer ho/bi mutations were observed at *NtBRC1-T*, most mutations at target sites were chimeric, carrying three to six mutated types without a non-edited type. Moreover, an identical 20bp spacer sequence with different PAMs (TGG/CGG) was designed to target a region of four high similarity *PEAPOD* (*NtPPD*) genes in allotetraploid tobacco. Sequencing results showed that 87.5% to 95.8% of T0 plants were mutated at each *NtPPD1* and *NtPPD2* loci, and 41.7% to 66.7% of them carried ho/bi mutations (Fig. 1h), suggesting that the pDC45 construct allows for robust editing efficiencies at targeted sites in tobacco.

To better evaluate the multiplex genome editing capability of pDC45_dsg systems, we selected two reported gRNAs, separately targeted to tobacco *fatty acid desaturase2* (*FAD2*) and *chloride channel* (*CLC*) alleles [4, 7], with extremely low ho/bi mutations (11.1% and 2.4%), and introduced them into tobacco using pDC45_dsg construct. Among 23 T0 plants, 65.2% and 34.8% of lines contained ho/bi mutations at *FAD2-2* and *CLC* sites, respectively, indicating that the pDC45_dsg system improved editing activities at poor efficiency sgRNA sites. Moreover, we examined dual gRNAs simultaneously targeted to *NtBRC1* and *NtCCD8* genes, which are correlated to the outgrowth of axillary buds in tobacco. The editing efficiency was up to 100.0% at targeted sites except that in the *NtBRC1-T* locus, and the ho/bi mutations of T0 lines at four sites were 95.0%, 90.5%, 81.0%, and 38.1%. The 33.3% of tested lines contained ho/bi mutations at all targeted sites, showing more vigorous shoot buds than wild-type (Fig. 1h, Supplementary Fig. 2). We constructed another cassette, targeting *NtWOX1* and *NtPDS* using the pDC45_dsg cassette. Among 27 T0 lines, 85.2% to 96.3% of them carried targeted mutations at each site, and 66.7% of tested plants were albino, bladeless, or albino bladeless leaves, due to complete disruption of all targeted alleles (Fig. 1i).

To expand the application of pDC45 expression systems, pDC45_MLOsg and pDC45_MYC2sg+ MLOsg were introduced into tomato to create mutations at target sites. The ho/bi mutations of T0 lines were respectively 73.9% and 45.5% for single and both target sites in tomato (Fig. 1h), indicating that pDC45 systems enable engineering the gene mutations in tomato genome.

*Flowering Locus T* (*FT*) is the critical positive regulator of flowering in plants, which has been integrated into the expression vector in an independent transcription manner to promote early flowering in tobacco [17, 18]. We speculated that the Cas9-FT integrated protein could be cleaved by self-cleaving 2A peptide, such that Cas9 and FT5 could function in gene editing events and simultaneously triggering early flowering in dicots. The resulting construct, pDC45_Fast, was used to test compatibility and editing efficiency in tobacco and tomato (Fig. 2a). We designed three separate sgRNAs targeting *NtPDS* and *NtBRC1* in tobacco, and the *SlMYC2* locus in tomato. Although ho/bi mutations at both NtBRC1 alleles were slightly lower than in the pDC45 cassette (28.6% vs. 35.5%), the ho/bi mutations at each site were similar efficiency to the pDC45 construct. Mutations at *NtPDS* and *SlMYC2* sites with relatively high efficiencies were detected in tobacco and tomato (Fig. 2b, 2d). Additionally, early bolting and fruit setting were observed in pDC45_Fast transformed plants, and the time for first fruit setting was respectively shortened to 87 days and 92 days for tomato and tobacco (Fig. 2c), compared to the normal regeneration process (around 120 and 150 days). No editing events were detected at any tested potential off-target sites, implying that the pDC45_Fast system is highly specific in plants (Fig. 2e).

**Fig.2.**
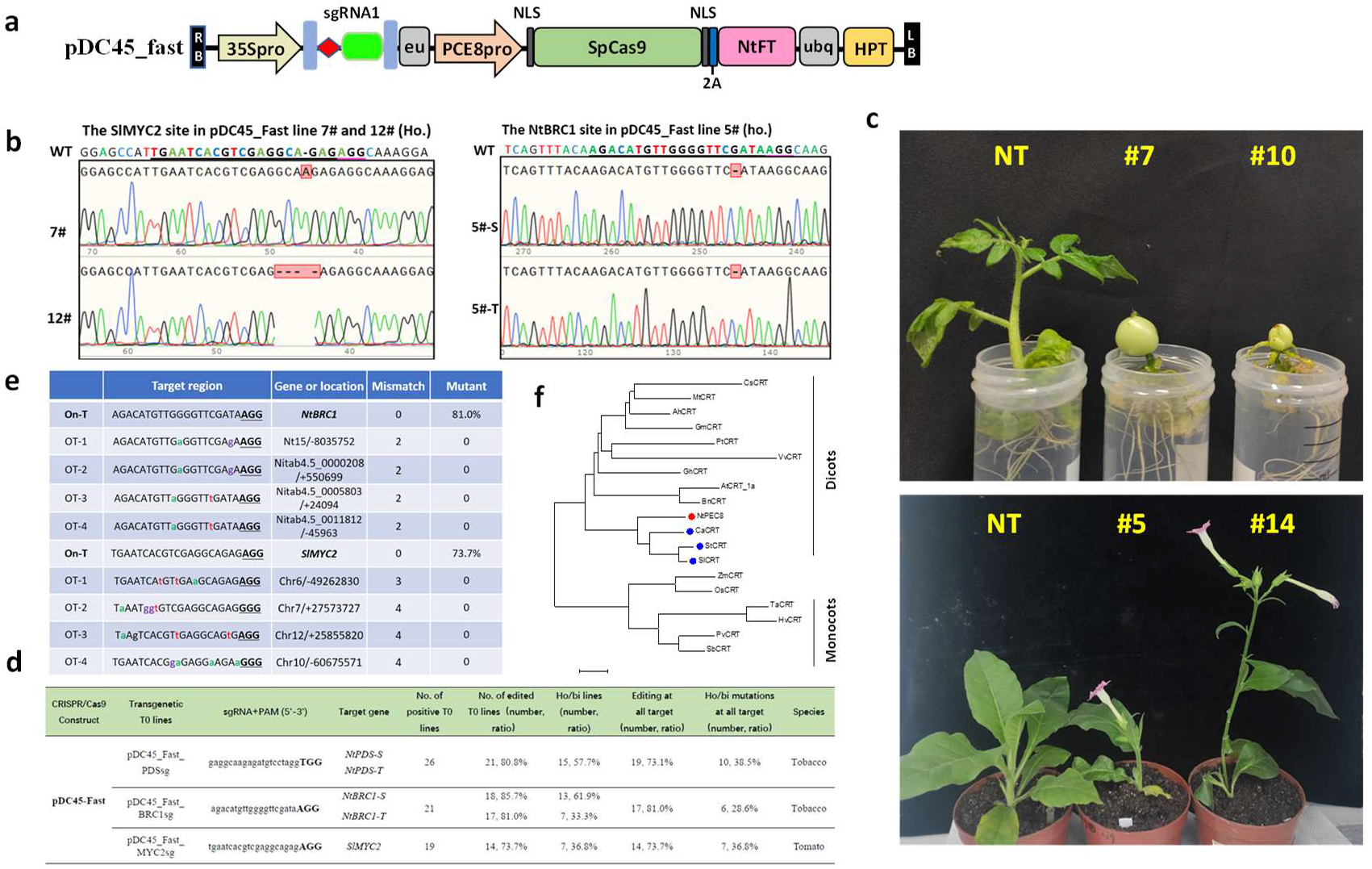
pDC_Fast mediated gene editing in dicot plants. **a** T-DNA structure of pDC_Fast vector, 2A, self-cleaving 2A peptide from porcine teschovirus-1; NtFT, Tobacco *Flowering locus T* gene. **b** Sequence chromatogram of PCR products of pDC45_Fast transgenic plants, the spacer and PAM sequence is underlined. **c** Early bolting and fruit setting in T0 transgenic tomato and tobacco. **d** Hi-TOM analysis of gene editing efficiencies of pDC_Fast transformed plants. **e** Detection of mutations at potential off-target sites in pDC45_Fast transgenic plants. **f** Phylogenetic tree of calreticulin-like proteins (CRTs) based on amino acid sequences from different species. Monocots and dicots were indicated in different groups. A filled circle with red and blue was marked before tobacco PCE8 and other CRTs in *Solanum* species.

Collectively, we identified a novel type promoter, *PCE8pro*, that can generate efficient gene editing in T0 generation plants. The pDC45 system, possessing *PCE8pro* and tandem tRNA-sgRNA units, is effective for multiplex gene editing in *Solanum*. Considering the high conservation of calreticulin-like protein (Fig. 2f, Supplementary Fig. 3), we anticipate that *PCE8*pro-controlled systems may be effective in other plant species.

## Conclusions

We developed the pDC45_dsg construct can generate more than 80.0% ho/bi mutations at *NtPDS* and *NtWOX1* sites in allotetraploid tobacco, and 52.2% of *NtPDS* ho/bi lines showed visible fragment deletions. We also observed a high multiplex genome editing capability of pDC45 construct in both tobacco and tomato, the editing efficiency ranged from 81.8% to 100% at each site. The pDC45_Fast system, can simultaneously fulfill gene editing and trigger early flowering, further shorten life cycle in T0 plants. The editing systems described here provide a new tool for improving targeted gene mutagenesis in dicots, and benefit future research in plants related to deciphering gene function, deletions of precursor miRNAs or chromosomal segments, and even large-scale gene editing.

## Methods

The primers used in this work are listed in Supplementary Table 1. The sequence of *PCE8pro, AtU6-sgRNA*, and *tRNA-BsaI_sg* is listed in the Supplemental material

### Plant material and sample preparation

The cultivated tobacco (*N. tabacum* cv. ‘honghuadajinyuan’) and Micro-Tom (*Solanum lycopersicum* cv.‘Micro-Tom’) were grown in the greenhouse. The genome DNAs of tobacco and tomato were extracted using the methods described previously [19].

### Vector construction

To construct p1300-U6_sg, the synthetic AtU6-sgRNA fragment (Supplemental material) was inserted into the *HindIII-EcoRI* site of modified pCAMBIA1300 binary vector (disruption of *BsaI* and *NheI* site). To increase the stability of Cas9 gene in plant, the Cas9-PA50 fragment was amplified with *Cas9-NcoI-F* and *Cas9-PA50-R* primer, then cloned into the *NcoI-BamHI* site of 18T-U6-sg-Cas9 vector [20], resulting in the generation of 18T-U6-sg-Cas9-PA50 vector. The AtUBQ1pro-Cas9-PA50 was subsequently introduced into the *HindIII-EcoRI* site of p1300-U6_sg vector, producing the pDC01 binary vector.

To construct the pDC30 and pDC40 vector, we respectively replaced the *AtUBQ1pro* fragment with *35Spro* and *PEC8pro* at *XmaI-NcoI* site of pDC01 vector. To obtain the pol II promoter-driven *tRNA-sgRNA* unit, the *2X35Spro* and *EU* terminator were amplified with corresponding primers (Supplementary Table 1), the *tRNA-BsaI_sg* was synthesized from BGI Tech (BEIJING LIUHE). Three fragments were integrated by overlap PCR methods, generating a 35Spro-tRNA_sgRNA-EU product. The 35Spro-tRNA_sgRNA-EU fragment with an adaptor, was amplified with *35S-CE-F* and *EU-CE-R* primer, and introduced into the *SpeI-SbfI* site of pDC40 via homologous recombination method, producing the pDC45 vector. The gRNAs used in this study were synthesized and inserted at *BsaI* site of pDC expressing vectors using T4 ligase (2011A, Takara). For dual sgRNA system, the fragment was amplified with the corresponding primers, and subsequently introduced into pDC45_dsg vector using Golden Gate Assembly Kit (E1601L, New England Biolabs). The tobacco FT gene was amplified using *NtFT-F* and *NtFT-R* primers, the *P2A-FT-F* and *NtFT-BamHI-CE-R* were used to generate FT-overlap fragment, and the Cas9-2A product was obtained using *CAS9-NcoI-CE-F* and *Cas9-linker-R* primer with pDC45 vector as a template. The Cas9-P2A-FT product with an adaptor, was generated by overlap PCR, and subsequently cloned into *NcoI-BamHI* site of pDC45 vector, resulting in the generation of pDC45_Fast. All constructions were confirmed by Sanger sequencing.

### Plants transformation

The stable transformation of tobacco was performed as previously described with minor modifications [19, 21]. Transgenic tomato was conducted using the Agrobacterium-mediated cotyledon transformation method as described previously [22].

### Mutation Detection

The genomic DNA was extracted from the tobacco regenerated lines using either Plant Direct PCR Kit (PD105-02, Vazyme) or NuClean Plant Genomic DNA kit (CW0531M, CWBIO) in accordance with relevant guidelines. The T0 plants were further confirmed by *SpCas9* amplification. To detect the targeted deletion and mutation of pDC45_PDSsg and pDC45_WOX1sg transformed lines, the PCR product amplified with PDS-del and WOX1-del primers (Supplemental table1), were either directly sequenced or ligated to the pEASY-Blunt Zero vector, and introduced into *E.coli* (Transgene, China). Ten or more colonies were sequenced to analyze the mutation types. To analyze the editing efficiency of diverse pDC systems in plants, the corresponding adaptor primers were used to amplify target fragments for deep sequencing. The data were collected and analyzed using the Hi-TOM platform [16].

### Off-target analysis

The potential off-target sites with 2 to 4 nucleotide mismatches compared to the sgRNAs of *NtBRC1* and *SlMYC2*, were searched using Cas-OFFinder tool [23]. Hi-TOM sequencing was conducted to detect the mutations of potential off-target sites in line 7 of tomato and line 5 of tobacco.

## Supporting information

Original data of fig.1h and 2d

## Acknowledgements

We thank Professor Jian-Kang Zhu for providing 18T-U6-sg-Cas9 vectors. This work was supported by The Agricultural Science and Technology Innovation Program (ASTIP-TRIC02), Central Public-interest Scientific Institution Basal Research Fund (1610232019001), China Tobacco Genome Project [110202001021 (JY-04)].

## Author contributions

C. D., Y. S., and Y-H. S. conceived and designed the experiments. B. L., Y. S., L. W., J. L., F. W., J. C., J. M., A. D., X. W., M. C., and C. D. performed the experiments and analyzed the data. C. D. and Y. S. wrote the manuscript. All authors discussed the results and approved the final manuscript.

## Supplementary Figure

**Supplementary Fig. 1.**
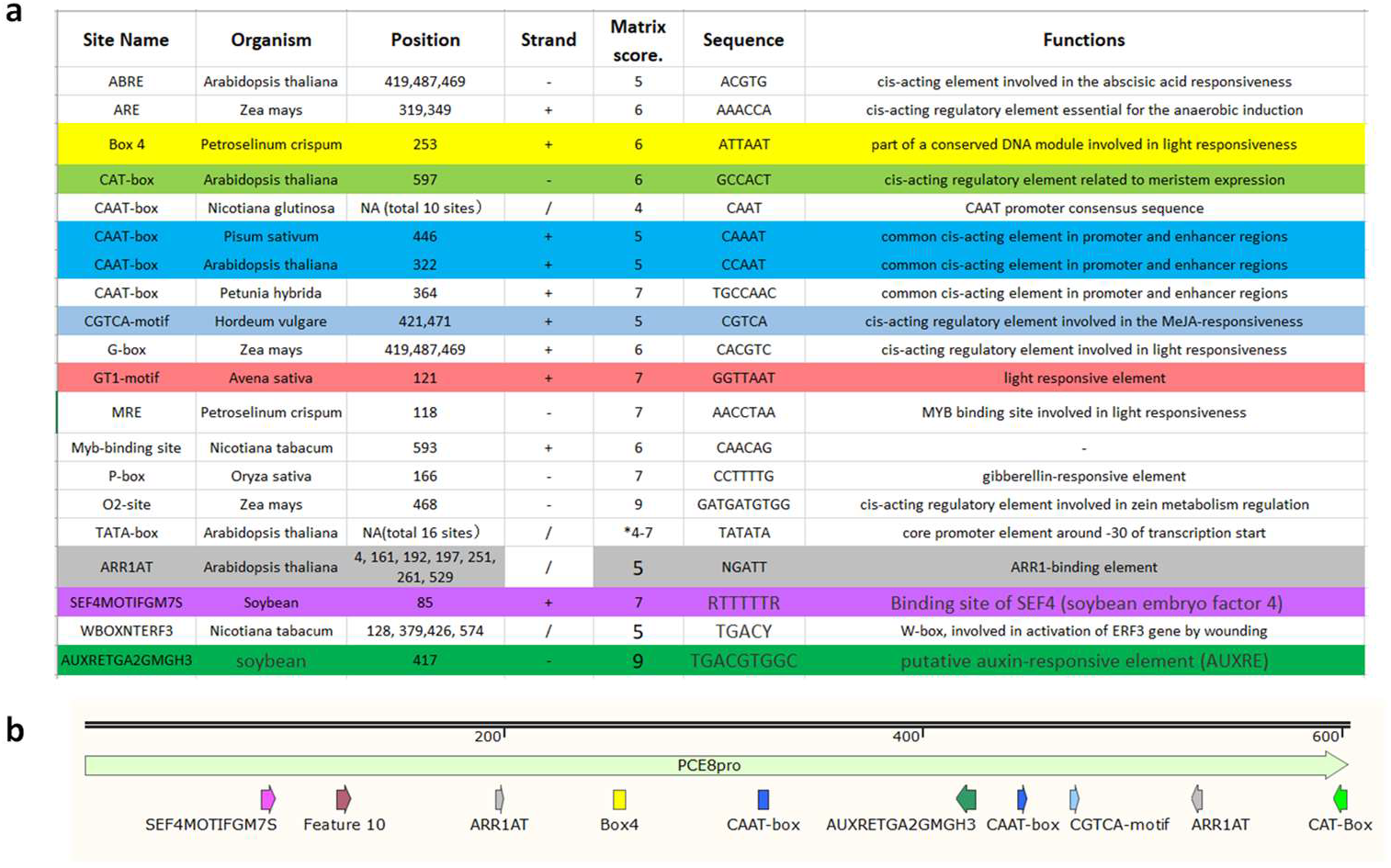
Putative cis-elements identified in the *<u>NtPCE8</u>* promoter sequence. **a** The core elements of *NtPCE8* promoter were analyzed using the online database, PlantCARE (http://bioinformatics.psb.ugent.be/webtools/plantcare/html/) and PLACE database (https://www.dna.affrc.go.jp/PLACE/?action=newplace). **b** A schematic of *NtPCE8* promoter with core elements. The colorful arrow and box shown the positions of functional elements in *NtPCE8* promoter.

**Supplementary Fig. 2.**
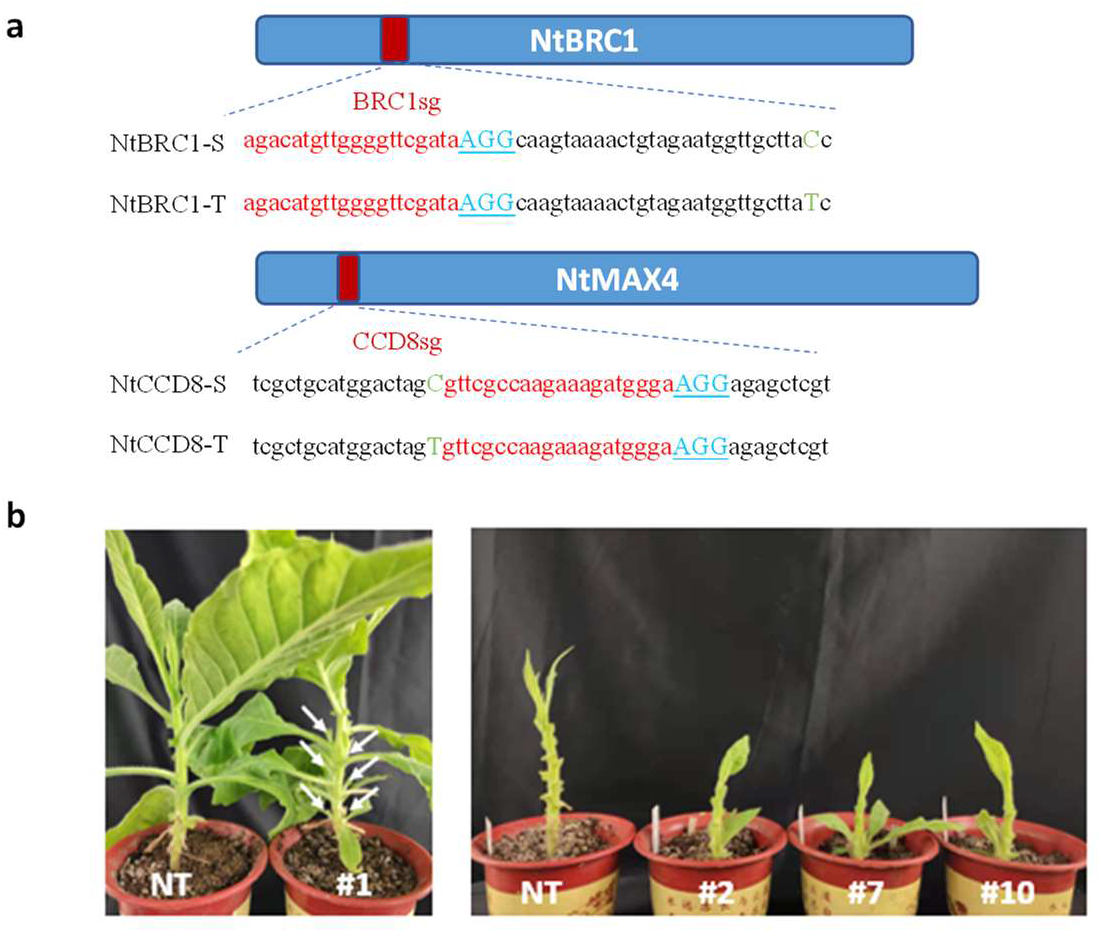
Gene editing at NtBRC1 and NtCCD8 site using pDC45_dsg systems. **a** The target position of NtBRC1 and NtCCD8 gene, red little letter show the space sequence, PAMs are underlined, green capital letter mean the SNP in tobacco S- and T-subgenome. **b** The phenotypes of T0 tobacco lines, white arrow show more shoot branching.

**Supplementary Fig. 3.**
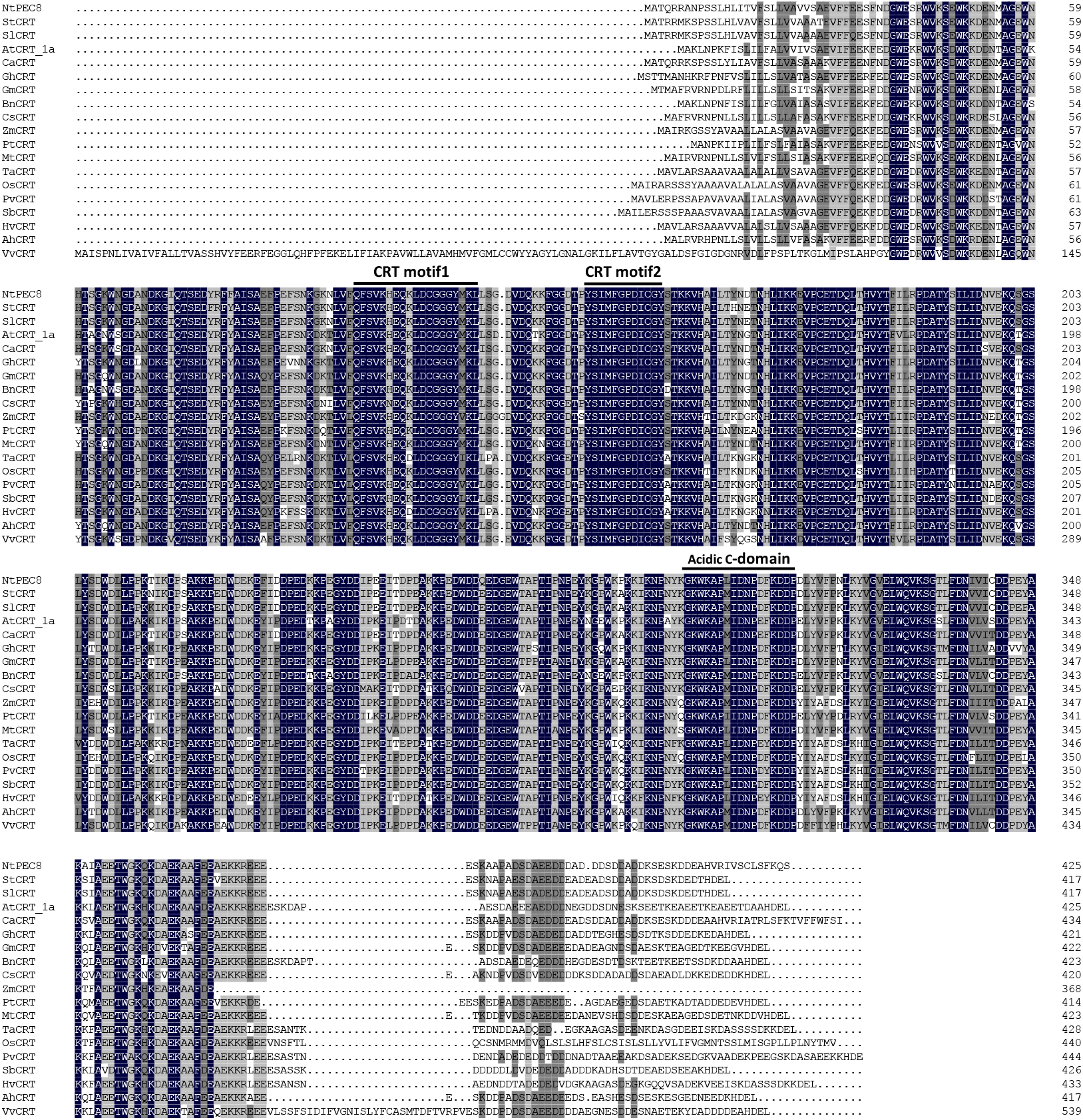
Sequence alignment of calreticulin-like (CRT) proteins in different plant species. Amino acid sequences of *CRT* genes were aligned using DNAMAN alignment program. Blackline indicates conserved CRT motif and acidic C-domain in plants. Cs, *Cucumis sativus* (XP_004144757.1); Mt *Medicago truncatula* (XP_024638651); Ah, *Arachis hypogaea* (XP_025674985); Gm, *Glycine max* (KAG5077294.1); Pt, *Populus trichocarpa* (XP_002318957.1); Vv *Vitis vinifera* (RVX01449); Gh, *Gossypium hirsutum* (XP_016667890.2); At, *Arabidopsis thaliana* (NP_176030.1); Bn, *Brassica napus* (XP_013687715); Ca, *Capsicum annuum* (PHT61563.1); St, *Solanum tuberosum* (KAH0766737.1); Sl, *Solanum lycopersicum* (XP_004230299.1); Zm, *Zea mays* (ONM52621.1); Os, *Oryza sativa* (KAF2922127); Ta, *Triticum aestivum* (XP_044391420.1); Hv, *Hordeum vulgare* (XP_044949696); Pv, *Panicum virgatum* (XP_039830141); Sb, *Sorghum bicolor* (XP_021307591).

## Supplemental material

### PCE8 promoter

AGTGG, cis-element related to meristem expression; GCCACGTCA, auxin-responsive element (AUXRE); TGACY, activation of ERF3 gene by wounding; CGTCA, MeJA-responsiveness

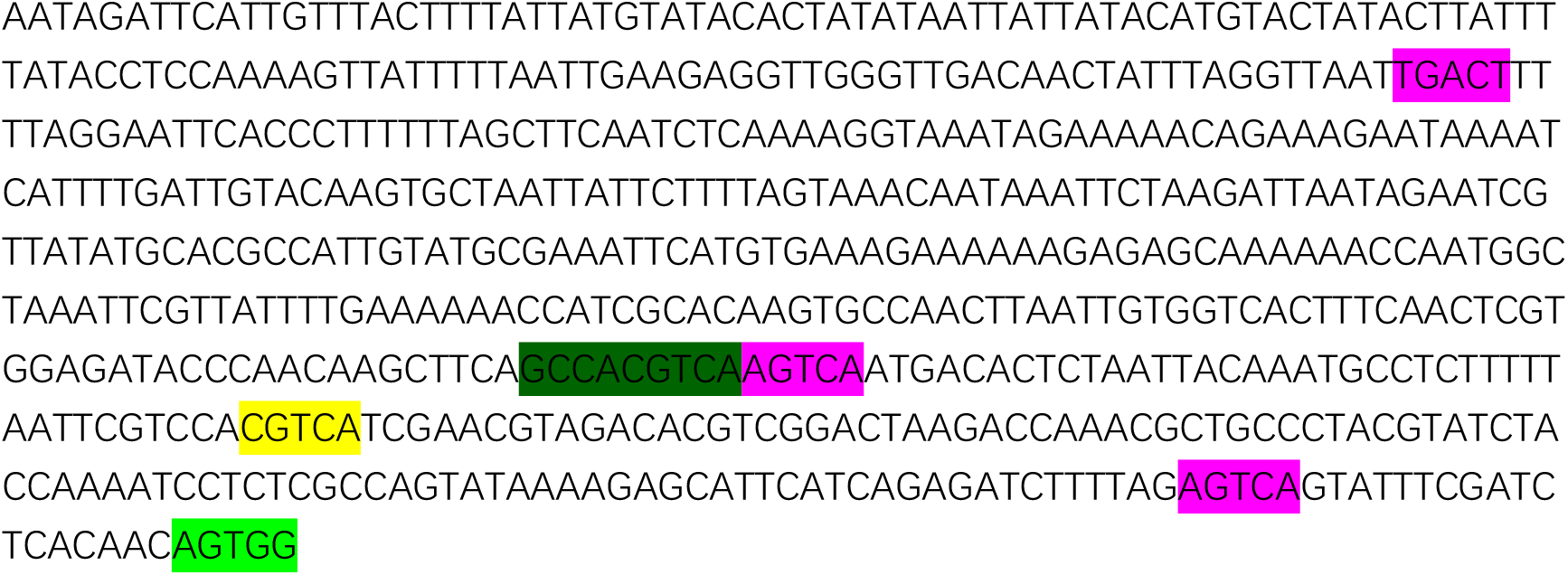

### AtU6-sgRNA

Light green, AtU6-26; Yellow, sgRNA scanfold

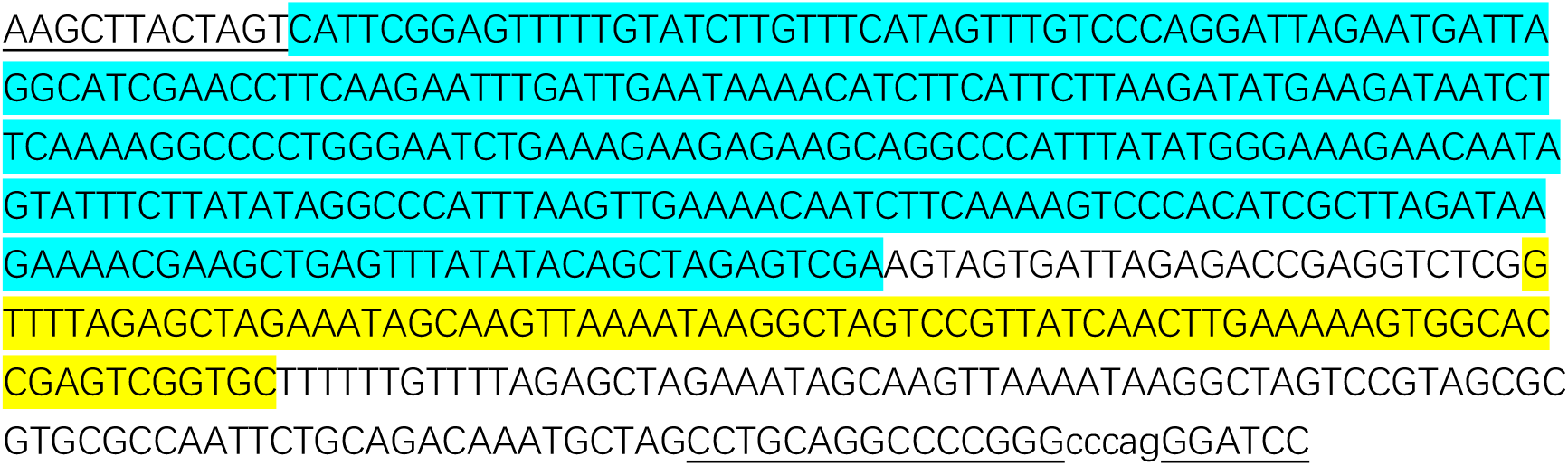

### tRNA-BsaI_sg

Green, tRNA; Yellow, sgRNA scaffold

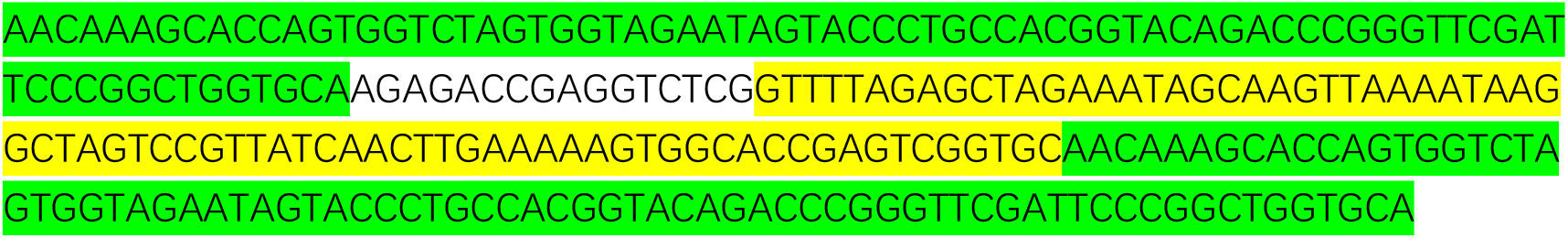

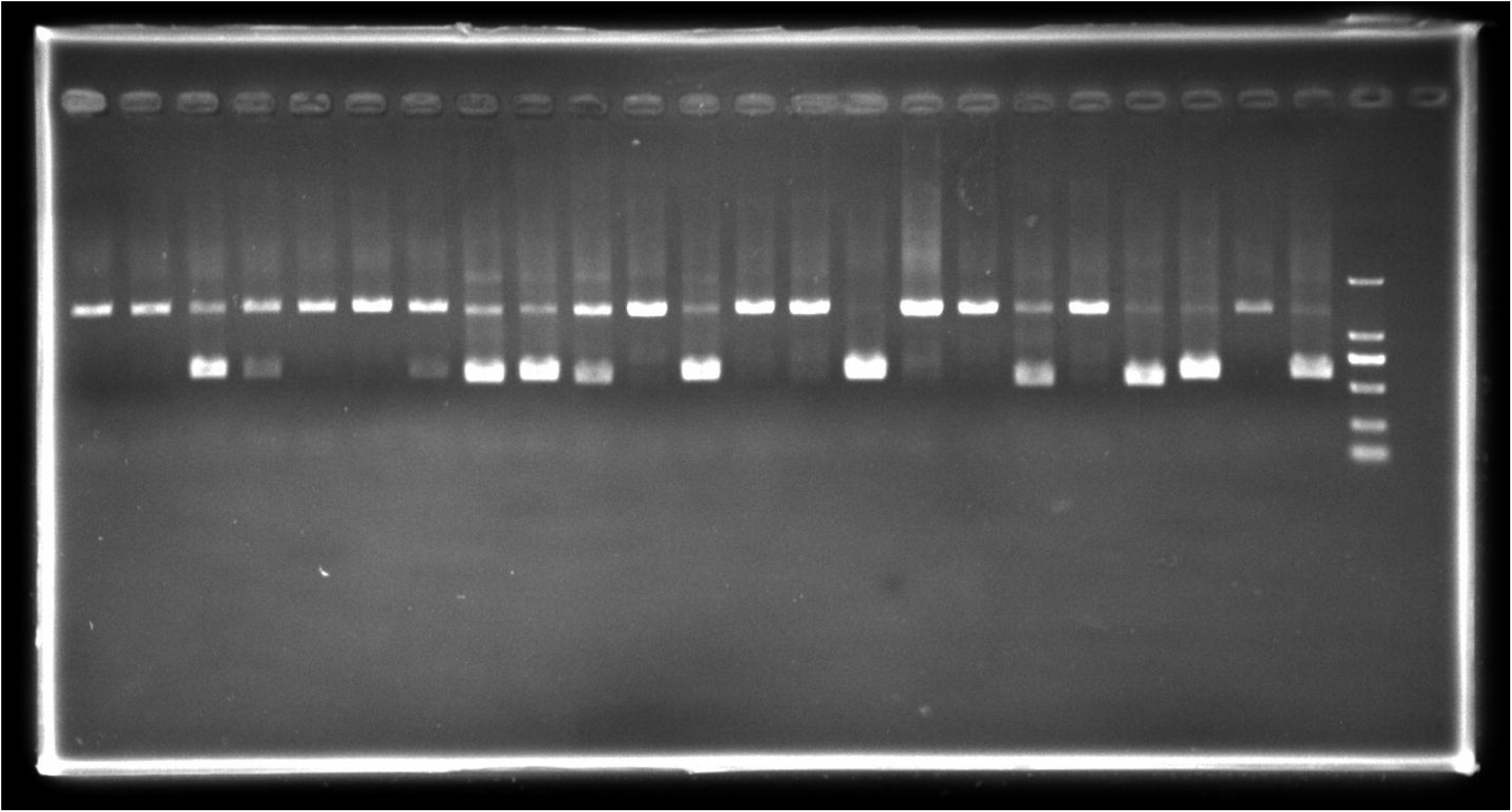

