## Supplementary material for "Highly efficient multiplex genome editing in dicots using improved CRISPR/Cas systems": Original data of fig.1h and 2d

| **CRISPR/Cas9Construct** | **Transgenetic  T0 lines** | **sgRNA+PAM (5’-3’)** | **Target gene** | **No. of positive T0 lines** | **No. of edited  T0 lines（number, ratio）** | **Ho/bi lines (number, ratio)** | **Editing at  all target（number, ratio）** | **Ho/bi mutations at all target（number, ratio）** | **Species** |
| --- | --- | --- | --- | --- | --- | --- | --- | --- | --- |
| **pDC30** | pDC30_PDSsg | gaggcaagagatgtcctagg**TGG** | *NtPDS-S* | 20 | 15, 75.0% | 7, 35.0% | 13, 65.0% | 5, 25.0% | Tobacco |
|  |  |  | *NtPDS-T* |  | 16, 80.0% | 10, 50.0% |  |  |  |
| **pDC45** | pDC45_WOX1sg | tgttggggatggccagttct**TGG** | *NtWOX1-S* | 29 | 24, 82.8% | 18, 62.1% | 24, 82.8% | 15, 51.7% | Tobacco |
|  |  |  | *NtWOX1-T* |  | 27, 93.1% | 23, 79.3% |  |  |  |
|  | pDC45_CENsg | ctcagtcacctctaaaccta**GGG** | *NtCEN2* | 22 | 21, 95.5% | 18, 81.8% | 20, 90.9% | 15, 68.2% | Tobacco |
|  |  |  | *NtCEN4* |  | 20, 90.9% | 15, 68.2% |  |  |  |
|  | pDC45_BRCsg | agacatgttggggttcgata**AGG** | *NtBRC1-S* | 31 | 29, 93.5% | 18, 58.1% | 28, 90.3% | 11, 35.5% | Tobacco |
|  |  |  | *NtBRC1-T* |  | 28, 90.3% | 11, 35.5% |  |  |  |
|  | pDC45_MLOsg | gagaaaatggagggcatggg**AGG** | *SlMLO* | 23 | 21, 91.3% | 17, 73.9% | 21, 91.3% | 17, 73.9% | Tomato |
|  | pDC45_PPDsg | gcctgtgatttattccatga**T/CGG** | *NtPPD1-S* | 24 | 21, 87.5% | 11, 45.8% | 18, 75.0% | 7, 29.2% | Tobacco |
|  |  |  | *NtPPD1-T* |  | 21, 87.5% | 11, 45.8% |  |  |  |
|  |  |  | *NtPPD2-S* |  | 22, 91.7% | 10, 41.7% |  |  |  |
|  |  |  | *NtPPD2-T* |  | 23, 95.8% | 16, 66.7% |  |  |  |
| **pDC45_dsg** | pDC45_PDSdsg | gctgcatggaaagatgatga**TGG** | *NtPDS-S/T* | 37 | 35, 94.6% | 33, 89.2% | 35, 94.6% | 32, 86.5% | Tobacco |
|  |  | gaggcaagagatgtcctagg**TGG** |  |  | 36, 97.3% | 34, 91.9% |  |  |  |
|  | pDC45_WOX1dsg | ggtttcaaaatcacaaggct**AGG** | *NtWOX1-S/T* | 28 | 26, 92.9% | 23, 82.1% | 25, 89.3% | 23, 82.1% | Tobacco |
|  |  | tgttggggatggccagttct**TGG** |  |  | 27, 96.4% | 24, 85.7% |  |  |  |
|  | pDC45_FAD2-2sg+CLCsg | ctgaaagcagtgaggtggga**TGG** | *NtFAD2-2-S* | 23 | 20, 87.0% | 15, 65.2% | 15, 65.2% | 6, 26.1% | Tobacco |
|  |  |  | *NtFAD2-2-T* |  | 21, 91.3% | 17, 73.9% |  |  |  |
|  |  | gcaccagcagctgcagggtc**AGG** | *NtCLC-S* |  | 16, 69.6% | 8, 34.8% |  |  |  |
|  |  |  | *NtCLC-T* |  | 19, 82.6% | 9, 39.1% |  |  |  |
|  | pDC45_CCD8sg+  BRC1sg | gttcgccaagaaagatggga**AGG** | *NtCCD8A* | 21 | 21, 100.0% | 20, 95.2% | 19, 90.5% | 7, 33.3% | Tobacco |
|  |  |  | *NtCCD8B* |  | 21, 100.0% | 19, 90.5% |  |  |  |
|  |  | agacatgttggggttcgata**AGG** | *NtBRC1-S* |  | 21, 100.0% | 17, 81.0% |  |  |  |
|  |  |  | *NtBRC1-T* |  | 19, 90.5% | 8, 38.1% |  |  |  |
|  | pDC45_WOX1sg+PDSsg | tgttggggatggccagttct**TGG** | *NtWOX1-S* | 27 | 23, 85.2% | 19, 70.4% | 23, 85.2% | 18, 66.7% | Tobacco |
|  |  |  | *NtWOX1-T* |  | 25, 92.6% | 23, 85.2% |  |  |  |
|  |  | gaggcaagagatgtcctagg**TGG** | *NtPDS-S* |  | 25, 92.6% | 20, 74.1% |  |  |  |
|  |  |  | *NtPDS-T* |  | 26, 96.3% | 23, 85.2% |  |  |  |
|  | pDC45_MYC2sg+MLOsg | tgaatcacgtcgaggcagag**AGG** | *SlMYC2* | 22 | 18, 81.8% | 12, 54.5% | 17, 77.3% | 10, 45.5% | Tomato |
|  |  | gagaaaatggagggcatggg**AGG** | *SlMLO* |  | 19, 86.4% | 16, 72.7% |  |  |  |

| **CRISPR/Cas9Construct** | **Transgenetic  T0 lines** | **sgRNA+PAM (5’-3’)** | **Target gene** | **No. of positive T0 lines** | **No. of edited  T0 lines（number, ratio）** | **Ho/bi lines (number, ratio)** | **Editing at  all target（number, ratio）** | **Ho/bi mutations at all target（number, ratio）** | **Species** |
| --- | --- | --- | --- | --- | --- | --- | --- | --- | --- |
| **pDC45** | pDC45_Fast_  PDSsg | gaggcaagagatgtcctagg**TGG** | *NtPDS-S*  *NtPDS-T* | 26 | 21, 80.8% | 15, 57.7% | 19, 73.1% | 10, 38.5% | Tobacco |
|  | pDC45_Fast_  BRC1sg | agacatgttggggttcgata**AGG** | *NtBRC1-S* | 21 | 18, 85.7% | 13, 61.9% | 17, 81.0% | 6, 28.6% | Tobacco |
|  |  |  | *NtBRC1-T* |  | 17, 81.0% | 7, 33.3% |  |  |  |
|  | pDC45_Fast_  MYC2sg | tgaatcacgtcgaggcagag**AGG** | *SlMYC2* | 19 | 14, 73.7% | 7, 36.8% | 14, 73.7% | 7, 36.8% | Tomato |
